# Characterisation of spinal ligaments in the embryonic chick

**DOI:** 10.1101/2025.04.22.650009

**Authors:** Sarah Hennigan, Ebru Talak Basturkmen, Rebecca A. Rolfe

## Abstract

Ligaments are important connective tissues within the musculoskeletal system that connect bone to bone and provide support and stability. The spine contains a number of ligaments that predominantly function in mechanical stabilisation and allow for certain ranges of spinal motion. Establishment of mechanical stability provided by spinal ligaments has not been described, and it is not known to what extent failure or inadequate spinal ligaments contribute to spinal conditions, such as scoliosis. While there are many similarities between ligaments and tendons, there is no experimental evidence investigating the development of these stability bearing tissues. This study uses the embryonic chick model *Gallus gallus* and investigates the development of spinal ligaments in the thoracic spine, examining structure and molecular expression across development. Findings show organisational changes in spinal ligaments in association with vertebral shape changes from cranial to caudal, with the anatomical identification of six vertebral ligaments in the thoracic spine. As development proceeds the size of the anterior longitudinal ligament, on the ventral surface of the vertebral body, and the supraspinous ligament, on the dorsal side of the spine, becomes greater, with the orientation of collagen fibres in the supraspinous ligament becoming more aligned. In addition, this study demonstrates that cell density decreases and nuclei become smaller and more circular across development. This study provides evidence that the embryonic chick is an appropriate model to study spinal ligament development and has added knowledge on the structural hallmarks of embryonic vertebral ligament tissues. These findings allow for subsequent investigation of the mechanical and molecular characteristics of spinal ligament development, for example useful for determining if *in utero* movement is important for the establishment of spinal ligament stability. Use of this model and integration of findings with additional models will provide knowledge of the contribution of spinal ligaments in spinal failure conditions.

## 1 Introduction

A healthy musculoskeletal system relies on interdependence between its component parts for functional success. Tendons and ligaments are closely related dense fibrous connective tissues that are key components of the system, playing vital roles in mobility and stability. While tendons connect muscle to bone to facilitate movement by transmitting load, ligaments join bone to bone and function to stabilise joints and guide movement through a normal range of motion(Bobzin et al., 2021). Ligaments are located throughout the body providing structural stability, most prominently in limb joints but also within the spine(Bogduk, 2016; Butt et al., 2015b). The principle functions of ligaments in the spine, spinal ligaments, are to hold the bony vertebrae together, stabilise the spine, protect the spinal cord and allow for limited physiologic ranges of motion(Bogduk, 2016; Butt et al., 2015b). Both tendons and ligaments rely on their strong and flexible structure of collagen fibrils, which are organised and bundled into sheaths(Benjamin and Ralphs, 1997; Bobzin et al., 2021). There are multiple similarities between tendons and ligament tissues, including protein and extracellular matrix composition, gene expression profiles and morphological properties, yet to date greatest attention has been paid to tendon tissues and as such are better-understood(Bobzin et al., 2021). There is increasing evidence surrounding the patterning, differentiation, and maintenance of anatomically distinct populations of tendons(Asahara et al., 2017), which highlights that much less is known about the specific structural, molecular and biomechanical mechanisms required for ligament development and homeostasis(Scaal, 2016), and nothing about spinal ligaments.

Much of our knowledge about spinal ligament structure and function comes from cadaveric studies providing basic anatomical descriptions(Butt et al., 2015b; Jiang et al., 1994; Panjabi et al., 1991a, b) and simple biomechanical information(Ivancic et al., 2007; Mattucci et al., 2012; Yoganandan et al., 2000; Zheng et al., 2018). It is well recognised though that more quantitative data on spinal ligament anatomy, studies on basic physical properties and investigations into the basic mechanisms through which ligaments adapt are required(Oxland, 2016). The major ligaments that stabilise the human spine are the supraspinous ligament (lies dorsally above the spinous process connecting the dorsal portion of each vertebrae, running from cranium to sacrum), the interspinous ligament (segmental ligaments throughout the spine), intertransverse ligaments (connecting transverse processes of adjacent vertebrae, present where transverse processes are present (the thoracic and lumbar regions)), ligamenta flava (connect the laminae of the spine, running within the spinal column), anterior longitudinal ligament (very wide and strong ligament covering the anterior body of the vertebrae from cervical vertebra 1 to sacrum) and the posterior longitudinal ligament (narrow, thin ligament located within the spinal column that runs from cervical vertebra 1 to sacrum) (Butt et al., 2015a). These ligaments operate along the length of the spine to allow for extension and flexion of particular regions. Ligaments are naturally designed so that when they are subjected to different movements they will respond based on their morphological composition(Hukins et al., 1990; Zheng et al., 2018). This architectural characteristic demonstrates the important interdependence of structural composition for physiologic function. While research on spinal ligaments has primarily focused on humans, some comparisons with animals spines have been done, finding that midline ligaments are present in all animals investigated, while lateral ligaments, such as the intertransverse ligaments are found in bipeds, such as human and aves but not in quadrupeds, suggesting that these ligaments play a unique role in the mechanical challenge of the erect spinal posture(Jiang et al., 1995). This suggests that while the quadrupedal mouse is more closely related to humans than the avian *Gallus gallus*, the chick could potentially be a more suitable model organism for investigating the embryonic development of spinal ligaments. Currently there is no experimental evidence profiling the developmental progression of spinal ligaments in any model organism. This study is the first to profile developing spinal ligaments to provide insight into whether the embryonic chick is an appropriate experimental model for the study of spinal ligaments, that can provide a basis for further investigations into the molecular and biomechanical mechanisms required for ligament development.

This study uses microanatomical approaches to identify and profile spinal ligaments along the length of the spine and across development, inferring nomenclature from mammalian anatomies. It investigates structural tissue organisation, collagen fibre alignment, candidate gene expression profiles and changes in cellular profiles in ligaments across development.

## 2 Materials and Methods

### 2.1 Egg incubation

Fertilised eggs (Ross 308, supplied by Allenwood Broiler Breeders), were incubated at 37.7°C in a humidified incubator. Work on chick embryos does not require a licence from the Irish Ministry of Health under European Legislation (Directive 2010/63/EU), all work on chick embryos was approved by the Trinity Ethics committee. Following 3 days of incubation, 5mls of albumen was removed from each egg using an 18-gauge needle. Embryos were sacrificed between E14 (HH40) to E20 (HH46). The final stages examined include E14, E16 and E20. Each egg was placed on ice for at least 15 minutes prior to harvest. Euthanasia of cooled embryos was performed by decapitation. Embryos were placed in to ice cold Phosphate Buffered Saline (PBS). Spines were either micro-dissected for RNA extraction or fixed in 4% paraformaldehyde (PFA) in PBS at 4°C for histological analysis.

### 2.2 Histological tissue processing and analysis

Fixed spines were placed in 10% EDTA (pH 7.4) for at least 7 days at 4°C for decalcification and subsequently dehydrated through a graded series of ethanol (ETOH) (25%, 50%, 70%, for a minimum of 1x 1 hour washes). Spines were stored in 70% ETOH prior to processing for paraffin embedding. Sub-dissection of spine regions were performed, with the use of the ribs as identification points. Separation of the thoracic vertebra 1 and 2 (T1-T2) segment was made using a scalpel blade and two transverse cuts, between cervical vertebra 14 (C14) and T1 and between T2 and T3. Partial dorsal portions of the ribs were left on the T1-T2 vertebral segment for orientation purposes for paraffin embedding. Paraffin processing involved subsequent 2 x 1 hour washes in 100% ETOH, 2 x Histoclear (National Diagnostics) washes (40 mins-1 hour each (dependent on stage)) and a minimum of 5 x 1 hour paraffin washes prior to embedding in paraffin. A full series of cross sections (8µm) through the T1-T2, T3-T5 and T6-7 vertebral segments were performed for each specimen. Additionally a full series of longitudinal (8µm) sections through the thoracic region for T1-T2, T3-T5 and T6-7 vertebral segments were prepared for each normal development specimen. Paraffin sections were dewaxed, rehydrated, and stained for cartilage with 1% Alcian Blue (pH1) in 0.1M hydrochloric acid (5 minutes), followed by 0.5% Picrosirius Red (45 minutes) to stain collagen, or stained with Haematoxylin (Harris Sigman HHS32) and Eosin (Sigman HT110232) or further processed for protein localisation. Histological specimens were photographed using an Olympus DP72 camera and CellSens software (v1.6).

### 2.3 Picrosirius red staining and polarised light microscopy (PLM)

Collagen fibre alignment was examined in longitudinal sections of supraspinous ligaments across development (E14, E16 and E20). Collagen was stained with Picrosirius Red (0.5% Direct Red 80 (Sigma 365548) in saturated Picric acid solution (Sigma P6744)) for 45 minutes, and the samples were examined under polarised light (Olympus BX41TF Microscope fitted with polariser and analyser filters). Images were captured using Ocular Software. The sections were consistently oriented with the ligament long axis running left to right on the imaging screen and images were captured with the 20X objective, first under white light to image the ROI, followed by two polarised light images (as previously described in (Rolfe et al., 2024)). The polarised light images were selected by first setting the polariser filter to 0⁰ and rotating the analyser filter until the brightest birefringence pattern was visible (Image 1; angle 0⁰) and then moving the polariser filter to 45⁰ (Image 2; angle 45⁰). Using ImageJ, the two images were merged, converted to 8bit and a central portion of the ligament cropped to 440 x 440 pixels (4.4 pixels/ µm) for further analysis. Fibre alignment analysis was performed on each ROI using the Plugin OrientationJ tool in ImageJ (Rezakhaniha et al., 2012), specifically using “Distribution for Histogram” and “Analysis for Colour Map” functions. This produces a colour map of orientations and allows a histogram of local orientations to be generated. For all timepoints 3 ROIs were measured for each independent biological sample (chick embryo) across developmental time, the 3 ROIs were analysed from 5 biological replicates for each stage. Fibre alignment analysis data were represented as proportion of fibres within four orientation categories (+/− 10⁰, +/− 10-20⁰, +/− 20-30⁰, +/− 30-40⁰) across each group for comparison and statistical analysis. Statistical analysis was carried out by univariate ANOVA followed by Tukey post hoc tests. *p≤0.05, **p≤0.01, ***p≤0.001

### 2.4 Immunohistochemistry and Collagen detection

Heat-mediated antigen retrieval was performed on histological sections using 0.01 M sodium citrate (pH 8) for 20 mins at 90°C prior to block of non-specific binding. Plasmid DNA encoding enhanced green fluorescent protein (eGFP) fused to the collagen binding protein (CNA35-eGFP) was kindly provided by the laboratory of Maarten Merkx, processed and purified as previously described (Mohammadkhah et al., 2017). Detection was performed as previously described (Rolfe et al 2024). In brief, blocking was performed using 1% bovine serum albumin (BSA) in PBS for 1 hr at RT, incubation in binding protein was carried out at a dilution of 1:50 in blocking solution, overnight at 4⁰C. Following subsequent PBS washes mounting with DAPI nuclear stain was performed (Prolong Gold antifade reagent with DAPI, Life Technologies). Fluorescent specimens were photographed using an Olympus DP72 camera and CellSens software (v1.6).

### 2.5 Image analysis and quantification of morphological and cellular parameters

Measurements were taken of the spinous process at cross sections at the mid-point of each vertebra, stained with alcian blue and picrosirius red. Mid-point sections were determined by the shape of the transverse process, with the longest and most full being defined as centrally located, and the middle being equidistant from the absence of spinous processes in transition zones between vertebrae. Using ImageJ, measurements were taken of the spinous process from where the ligamentum flavum connects to the vertebral body to the dorsal tip of the spinous process. Width was measured perpendicular to (+/− 1°) and at the midpoint of this line. Length was divided by width to give aspect ratio (AR). Three technical replicates were taken per vertebra, with each vertebra being categorised as being T1-T2 (n=5), T3-T5 (n=5) or T6-T7 (n=6).

The width of anterior longitudinal ligaments and supraspinous ligaments were measured from midline longitudinal sections stained with alcian blue and picro sirius red. Midline longitudinal sections were identified by the morphological characteristics of the spinal cord and the spinous process. Length measurements, perpendicular to vertebral cartilage, were taken across three technical replicates for each biological replicate (n=5, for each E14, E16, E20). The supraspinous ligament adjacent to the spinous process at the cross-sectional midpoint of the vertebral body (identified using the method described above) was measured using the polygon selection tool. Outlines of one branch of the supraspinous ligament were made to assess the cross-sectional area and shape of the ligament. Three technical replicates were measured per biological replicate (E14 n=6, E16 n=7, E20 n=8).

Cell density was measured from supraspinous ligaments from midline longitudinal sections (identified as described above) stained with haematoxylin and eosin using the Cell Counter plugin for ImageJ. All nuclei were counted, including partially visible nuclei and edge nuclei. All analysis of densities represent a 10,000μm^2^ region of interest (ROI). Three technical replicates per biological replicate (E14 n=5, E16 n=4, E20 n=5) were measured. Nuclear shape and size were measured in supraspinous ligaments from midline longitudinal sections (identified as described above) stained with DAPI. Regions of interest of 50μm x 50μm were analysed as previously described (Rolfe et al., 2024). In brief, cropped images colour channels were split, images were binarised to distinguish individual nuclei, noise was removed using either the despeckle or freehand selection tools. Nuclei were analysed independently, with the total number of nuclei across three biological replicates for each stage (a minimum of 3 ROI for each stage) summing to n=440 (E14), n=564 (E16) and n=445 (E20).

### 2.6 Gene expression – RNA extraction, cDNA synthesis, qRT-PCR

The dorsal supraspinous ligament was sub dissected from the thoracic region between cervical vertebra 14 and lumbar vertebra 1. Ligaments from each embryo were placed in TRIzol (Thermo Fisher Scientific), stored at −70°C. Mechanical homogenisation was performed using polypropylene pestles and high speed vortexing. RNA was extracted from the stored TRIzol tissue homogenate with chloroform and purified with a TRIzol Purelink kit (Invitrogen) following manufacturers’ instructions. Typically, mRNA was re-suspended in 30-50 µl of nuclease-free water and quantified using the Qubit 2.0 Quantitation System (Invitrogen). mRNA was reverse transcribed using a standard quantity of total RNA (500 ng) and High Capacity cDNA Reverse Transcription Kit (Applied Biosystems™: 4368814), diluted 1:5 with RNase free water. Chick primers for genes of interest (Table 1) were synthesised by Merck. *Gapdh* was used as the normaliser transcript. Real-time PCR quantification was performed using an ABI 7500 Sequence Detection system (Applied Biosystems) using SYBR green gene expression quantification (Applied biosystems). For each well 5 μl of cDNA preparation, 10 μl of 2x SYBR green PCR master mix (Applied Biosystem), 0.5 µl (10 μM) of each primer and 4 µl RNase free water was prepared. Samples were assayed in triplicate in one run (40 cycles), which was composed of three stages, 95°C for 10 min, 95°C for 15 sec for each cycle (denaturation) and 60°C for 1 min (annealing and extension). Data were analysed using relative quantification and the Ct method as described previously (Livak and Schmittgen, 2001). The level of gene expression was calculated by subtracting the averaged Ct values (Ct is the threshold cycle) for *Gapdh* from those of the gene of interest. The relative expression was calculated as the difference (ΔΔCt) between the Ct of the test stage and that of the compared stage. The relative expression of genes of interest were calculated and expressed as 2-ΔΔCt. Relative quantification values are presented as fold changes plus/minus the standard error of the mean relative to the comparator group, which are normalised to one.

**Table 1:**
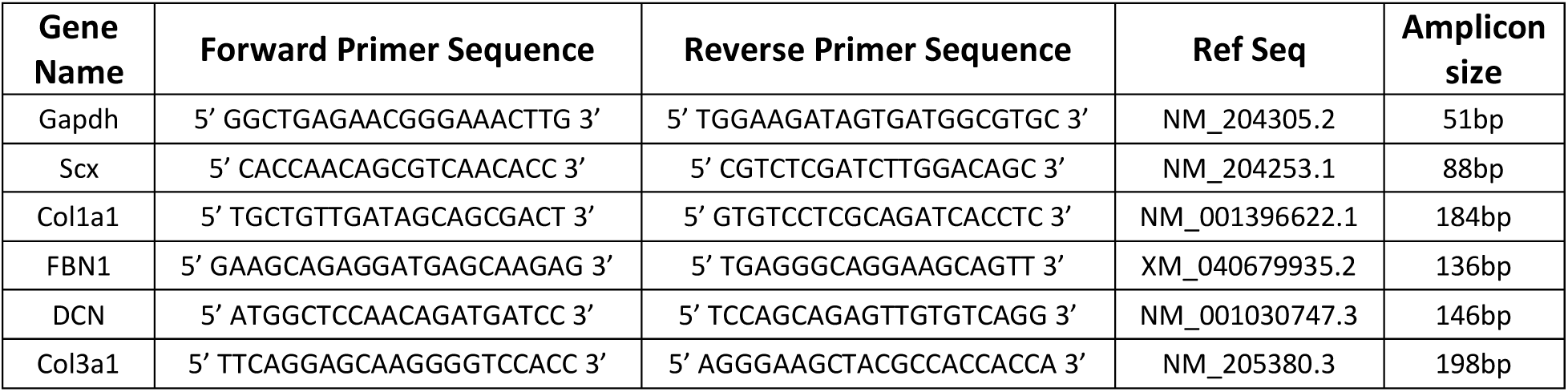
Details of primers used in qRT-PCR.

### 2.7 Statistical analysis

A range of 5 – 8 independent biological replicates (embryos) were analysed for all normal development stages (E14, E16 and E20). For morphological analysis in the transverse plane n=6 for E14, n=7 for E16 and n=8 for E20 were measured. In the longitudinal plane n=5 biological replicates for all stages were assessed, which were used for PLM analysis, measuring the width of both ALL and SSL, and cell density analysis. For transcript expression analysis n=6 for E14, n=5 for E16 and n=7 for E20 independent biological replicates were measured. For nuclear analysis n=3 for E14, E16 and E20 biological replicates were assessed with 440 nuclei from E14, 564 nuclei from E16 and 445 nuclei from E20 replicates analysed for nuclear shape. For statistical analysis SPSS statistics (IBM^®^, v.27) or R version 4.4.1 (© 2024 The R Foundation for Statistical Computing) and RStudio version 2024.12.1 Build 563 (© 2009-2025 Posit Software, PBC) were used. All data presented are mean plus/minus standard error of the mean (SEM). Significance was determined by one-way ANOVA, followed by Tukey’s post hoc test with a 95% confidence interval. P-values of ≤ 0.05 were considered significant.

## 3 Results

### 3.1 Spinal ligament anatomy in the embryonic chick is homologous to that of the human spine, while morphology varies based on orientation of visualisation

The morphological identification of six independent spinal ligaments in the embryonic chick spine was achieved in this study. The latest stage of embryonic development, E20, was examined in detail to reveal the six ligaments positioned adjacent to developing cartilaginous vertebrae with distribution in locations homologous to those in the mammalian spine (Figure 1). Using specific morphological features of developing cartilaginous vertebrae in conjunction with histological staining in two orientations it was possible to identify spinal ligaments in both the transverse (Figure 1A) and longitudinal plane (Figure 1B). Taking advantage of picrosirius red staining of collagenous tissues in combination with anatomical vertebral morphological features, six spinal ligaments were identified. These include: the anterior and posterior longitudinal ligaments (all and pll) (Figure 1A i & ii) positioned ventrally and dorsally to the vertebral body (vb), the ligamentum flavum (lf) (Figure 1A iv) in the vertebral canal dorsal to the spinal cord (sc) and ventral to the cartilaginous spinal process and the supraspinous ligament (ssl) dorsal to the spinous process (Figure 1A v). Due to their discrete positions the intertransverse ligaments (itl) were identified only in the transverse plane adjacent to transverse processes (tp) (Figure 1A iii) and the interspinous ligaments in the longitudinal plane nestled between the cartilaginous spinous process (Figure 1B iii). Individual collagen fibres were visualised within all of the spinal ligaments, with the arrangement of each ligament exhibiting unique features. The anterior longitudinal ligament (all) covers the entire ventral surface of the vertebral body, with apparent variability in thickness dependent on the vertebral body morphology as visualised in the longitudinal plane (Figure 1B), while the posterior longitudinal ligament (pll) is located in the vertebral canal.

**Figure 1:**
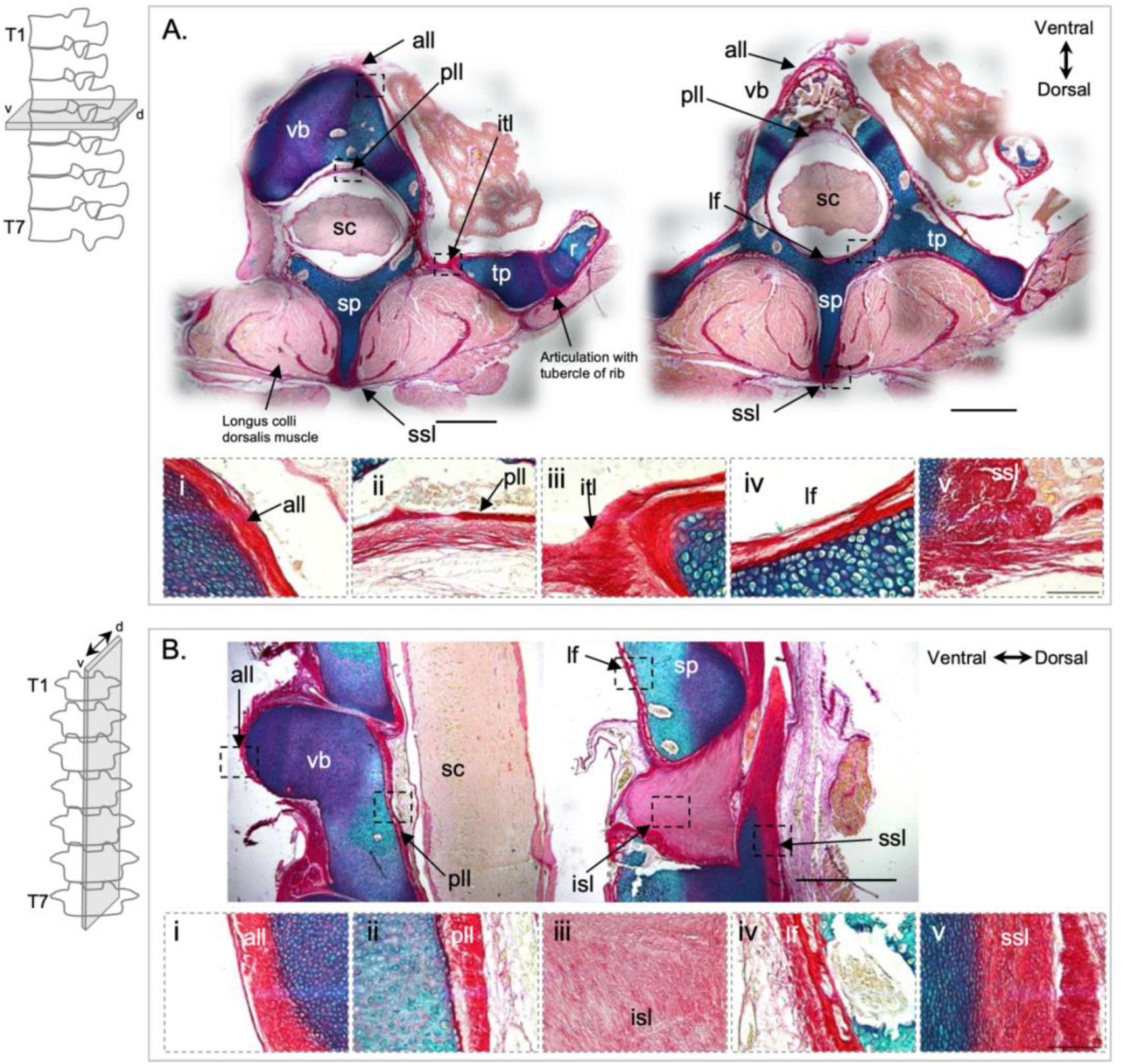
Spinal ligaments in the embryonic chick are homologous to those in the human spine. Histological visualisation of spinal ligaments in the thoracic (T) region are shown, with a schematic demonstrating orientation for each with line diagrams. Representative cross (A) and longitudinal (B) sections of the thoracic region of chick spines at embryonic day 20 (E20) stained with alcian blue, staining cartilaginous vertebrae and picrosirius red, staining collagenous tissues, ligaments and muscle. Two cross sectional views are shown (A) and one midline longitudinal section (B) with morphological vertebral features labelled including the ventral vertebral body (vb), dorsal spinous process (sp) and lateral transverse processes (tp). Higher magnification boxed regions are shown for all ligaments identified in both orientations, these include the anterior longitudinal ligament (all) (i), posterior longitudinal ligament (pll) (ii), intertransverse ligament (itl) (A iii), interspinous ligament (isl) (B iii), ligamentum flavum (lf) (iv) and supraspinous ligament (ssl) (v). Spinal cord (sc). Scale bars 1cm (A-B), 100μm (i-v). v: ventral, d: dorsal.

In order to understand morphological changes in ligament structure across development it was important to ensure that precise anatomical sites and orientations were compared. Here we profiled cross sectional representative views of vertebrae and ligaments in the cervical and thoracic regions, with a sharper focus on multiple levels within the thoracic region (Figure 2). This analysis was inspired from observations of large natural variations in cartilaginous vertebral morphologies and collagenous ligament patterns in the two dimensional transverse orientation from cranial to caudal (Figure 2). With depth the cross sectional morphology of embryonic cartilaginous vertebral bodies and spinous processes change, not only from the cervical to the thoracic region, but also within each region, as shown for three representative levels of thoracic vertebrae in the transverse plane (Figure 2). The ventral surface of vertebrae in the cervical region have a short central pointed shape (Figure 2A), while in the thoracic region the upper most vertebrae have three ventral projections with concave indentations between them (Figure 2B), while the mid thoracic vertebrae (T3-T5) demonstrate a triangular shaped vertebral body with a prominent midline ventral projection (Figure 2C) and the lower thoracic (T6-T7) vertebrae having a ventral convex vertebral body (Figure 2D). The dorsal spinous process remains present with depth, with an elongation and thinning at thoracic levels T3-T5 and T6-T7 compared to T1-T2 (Figure 2 C-D, Supplementary Figure 1B). The anterior longitudinal ligament (all, indicated by black arrowheads in Figure 2) conforms to the shape of the ventral vertebral body, while the dorsal supraspinous ligament (ssl, indicated by green arrowheads in Figure 2) displays a symmetrical pattern on both lateral sides of the spinous process, present in one bundle in the cervical region and two bundles throughout the thoracic regions, with multiple branching bundles evident within the dorsalis muscle between T3 and T7 (Figure 2B-D). The elaborate patterns of branching of the supraspinous ligament and changing shape profiles of the morphological features of vertebrae are depicted in supplementary figure 1, with an approximate distance of 1800μm represented for the images shown from top to bottom.

**Figure 2:**
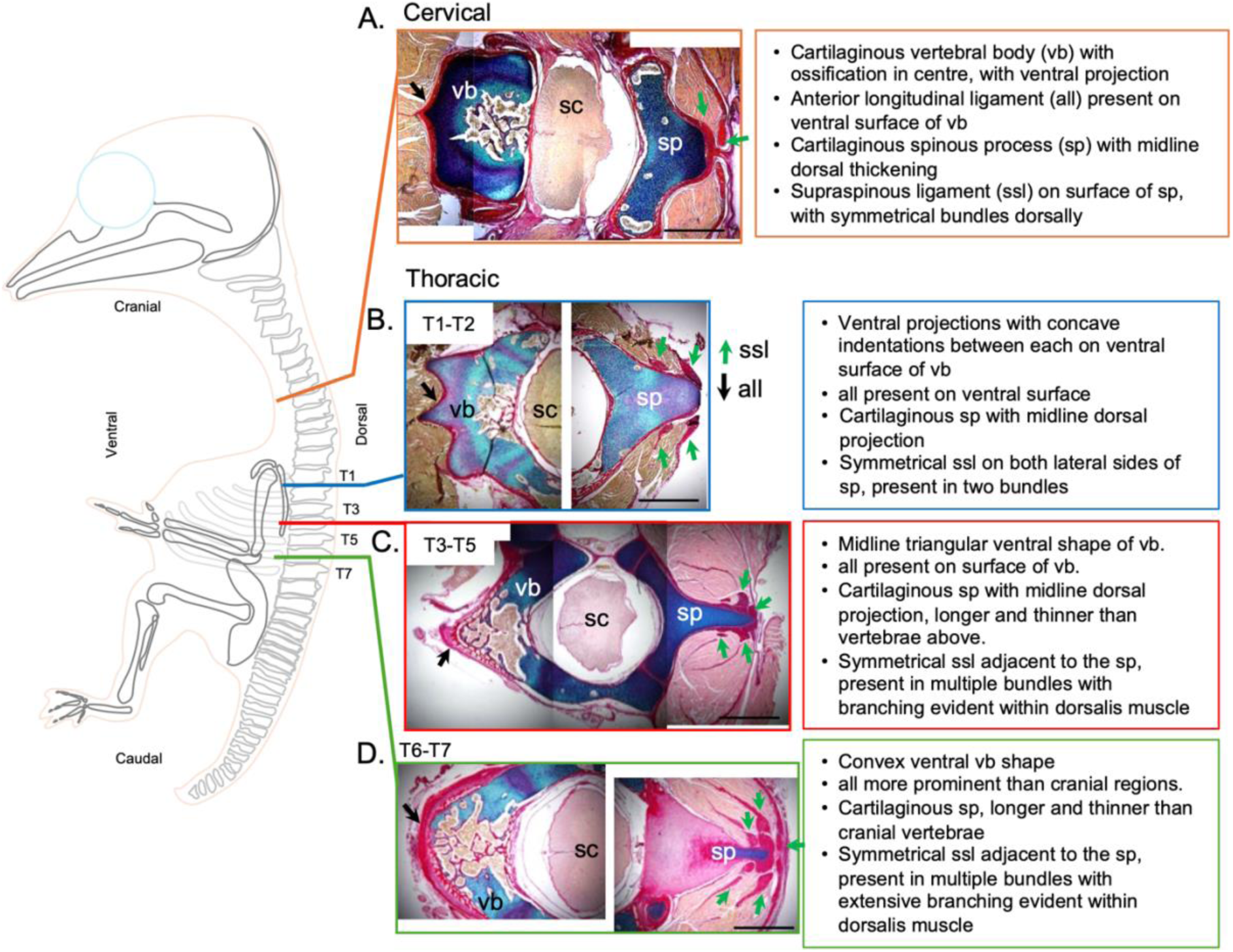
Morphological variation in vertebral shape and associated spinal ligaments in transverse orientations from cranial to caudal in the embryonic chick. Schematic lateral view of embryonic chick skeleton and histological representative transverse sections of the cervical (A) and thoracic (T1-T2 (B), T3-T5 (C), T6-T7 (D)) regions at embryonic day 20 (E20) stained with alcian blue and picrosirius red. Morphological features labelled include the cartilaginous ventral vertebral body (vb), dorsal spinous process (sp) and spinal cord within (sc). The anterior longitudinal ligament (all) and supraspinous ligament (ssl) (deep red) are indicated with black and green arrow heads respectively. Scale bars 1000μm.

### 3.2 The size, shape and molecular expression of spinal ligaments change during late stages of embryonic development

An investigation of spinal ligament morphological changes across development demonstrate that ligaments change in size and shape, from E14 to E20 (Figure 3), this corresponds to the maturation of vertebra, with obvious ossification occurring in vertebral bodies from E16 and progression at E20 (Figure 3B). Comparisons of changes in ligament cross sectional area and thickness were made across stages E14, E16, and E20 in the thoracic region from transverse and longitudinal sections respectively. Due to the elaborate transverse pattern of the supraspinous ligament with extensive branching demonstrated within the dorsalis muscle, assessment of the size of one of the upper ligament bundles was performed, with representative views of each bundle at each stage shown (Figure 3C). An increase in the cross sectional area of the upper bundle of the supraspinous ligament, measured on one side of the spinous process, was observed from E14 – E20 (10406.50±2129μm^2^ to 24260.58±4032μm^2^ p= 0.0195) (Figure 3C, D). Representative indications of thickness measurements of the anterior longitudinal ligament (all) and the supraspinous ligament (ssl) are indicated in figure 3E (yellow lines). The anterior longitudinal ligament (all) increases in thickness on the ventral surface of the vertebra between E14 to E20 (35.4±3.4μm to 58.4±2.8μm p=0.002) and E16 to E20 (44.7±4.3μm to 58.4±2.8μm p=0.044) (Figure 3 H), with representative views shown (Figure 3F). While the supraspinous ligament, measured at the most dorsal point of the cartilaginous spinous process (Figure 3E, yellow line) show a near doubling of thickness between both E14 to E20 and E16 to E20 (49.1±6.7μm to 85±5.9μm p=0.002 and 55.6±3.5μm to 85±5.9μm p=0.007, respectively) (Figure 3H).

**Figure 3:**
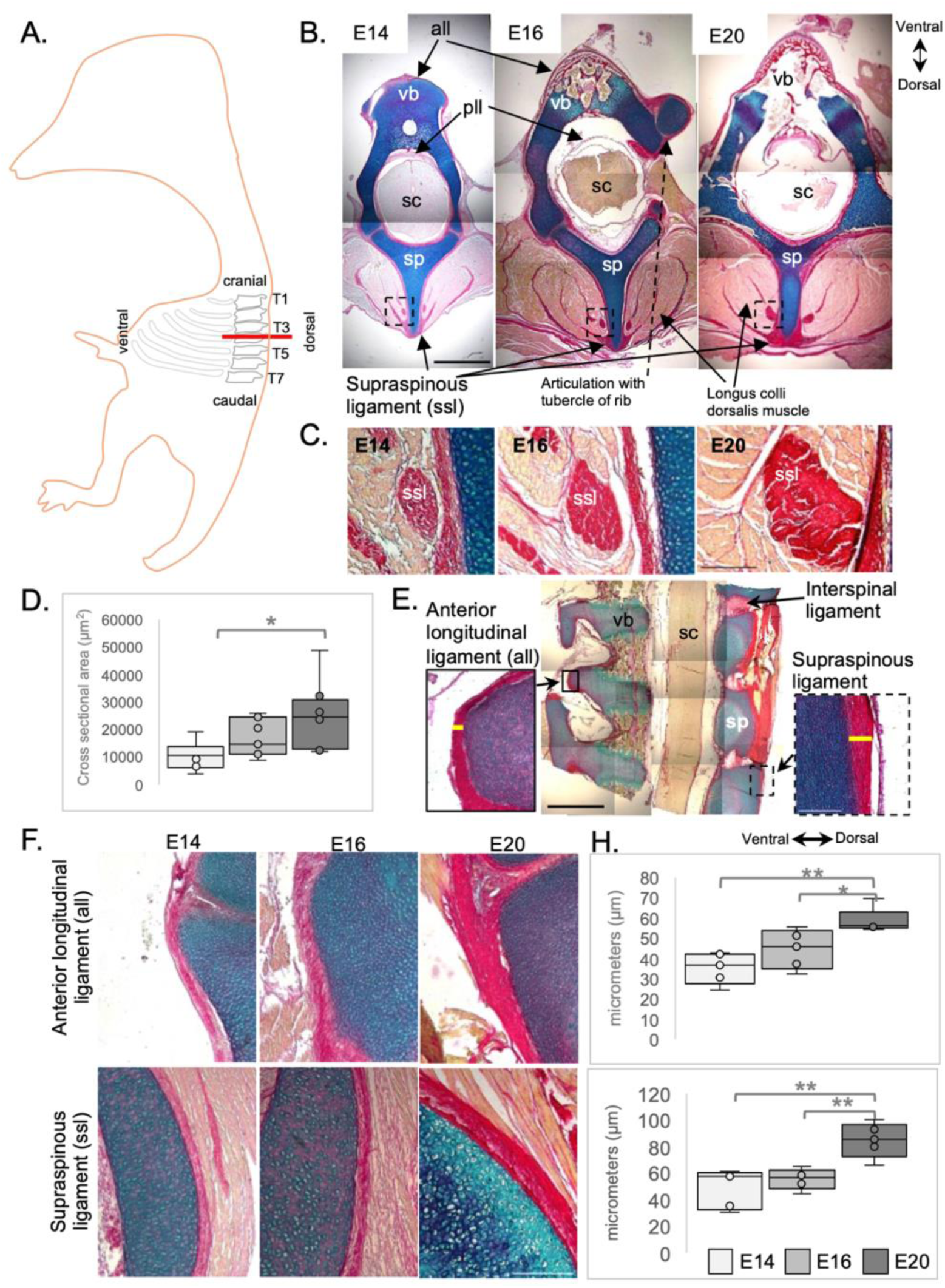
Spinal ligaments change in size and shape across development in the embryonic chick (E14-E20) A. Schematic lateral view of an embryonic chick highlighting the thoracic (T) region and the level of vertebrae depicted in B indicated with a red horizontal line. B. Representative transverse sections of thoracic vertebrae (between T3-T5) at E14, E16 and E20 showing cartilage (blue) in the vertebral bodies (vb) and spinous process (sp), collagen in ligaments (red) and muscle (light pink) surrounding the spinal cord (sc). C. Representative views of the upper bundle of the supraspinous ligament across development. D. Box plot showing change in the cross sectional area of the ssl across development. E. Longitudinal sections illustrating the width measurements of the anterior longitudinal ligament (all) and the supraspinous ligament (ssl) yellow lines. F. Representative views of all and ssl across development, demonstrating increases in width of both ligaments (H) across development. *p≤0.05, **p≤0.01, ***p≤0.001. Scale bar in (B, E) 1000µm, (C, E, F) 100µm.

Gene expression profiling of in vivo chick supraspinous ligaments from E14 to E20 show that the relative expression of *Scx* decreases between E14 and E16 (p=0.0002), and is further reduced at E20 by greater than 8 fold compared to E14 (p=0.000000006) (Figure 4). Decreases in collagen specific genes, *Col1a1* and *Col3a1* expression were observed between E16 to E20 (p=0.005 and p=0.017, respectively) (Figure 4). Maintenance of expression of the extracellular matrix glycoprotein gene *Fibrillin1* and proteoglycan gene *Decorin* were observed across the late stages of spinal ligament development.

**Figure 4:**
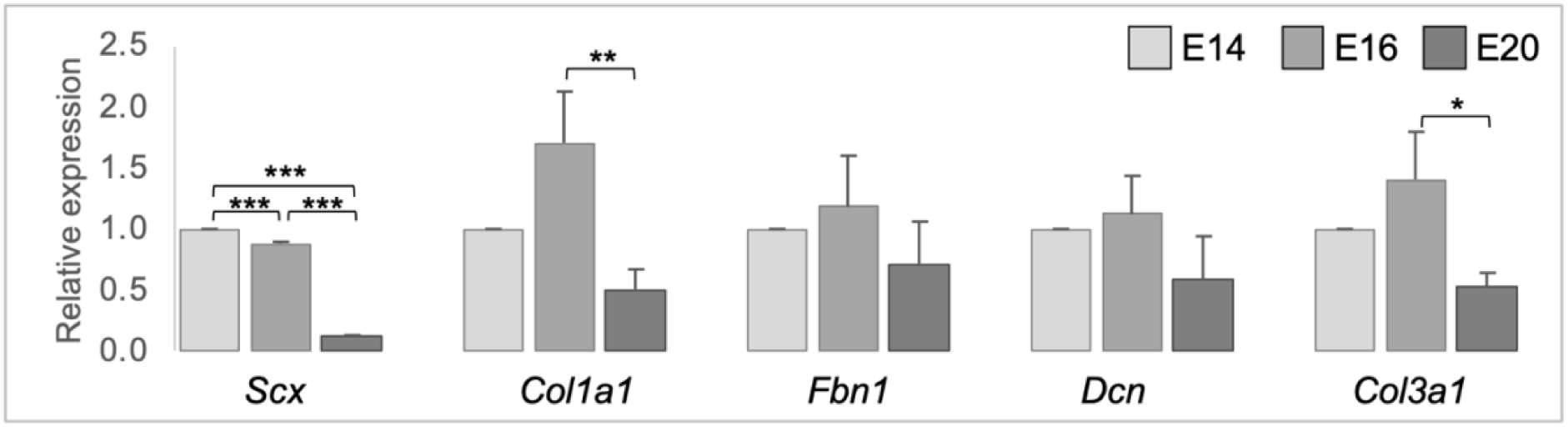
Gene expression analysis of micro-dissected dorsal supraspinous ligaments show a reduction in the expression of the transcriptional regulator Scx across development, reduced expression of collagen genes at E20 and the maintenance of expression of extracellular component genes fibrillin1 (*Fbn1*) and decorin (*Dcn*). *p≤0.05, **p≤0.01, ***p≤0.001.

### 3.3 Collagen fibre alignment increases in spinal ligaments across development

From the earliest stage examined (E14), collagen deposition was detected on all spinal ligaments visualised (Figure 3), with no apparent difference in collagen levels, through picrosirius staining, in the anterior longitudinal ligament or the supraspinous ligament across E14 to E20 (Figure 3F). To look closer at deposited collagen over time, picrosirius red staining (PSR) combined with polarised light microscopy (PLM) was used to analyse the spatial organisation of the collagen network (Liu et al., 2021a), as performed previously on embryonic tendons (Rolfe et al., 2024); in particular permitting quantitative analysis of collagen fibre alignment as well as reflecting the maturity of the deposited collagen (Patel et al., 2018). The colour of the birefringent light has been previously taken to indicate increasing fibre thickness and maturity from green to yellow, to orange and red. Analysis of collagen fibres in the supraspinous ligament shows progressively more alignment of fibres across late development (Figure 5), with a representative specimens for each stage demonstrated in figure 5A, and a full profile of all biological specimens in supplementary figure 2. Quantification of the proportion of fibres in orientation groups; +/− 10⁰, +/− 10⁰- 20⁰, +/− 20⁰- 30⁰ and +/− 30⁰- 40⁰ of the longitudinal axis across time, shows that the proportion of fibres in the +/− 10° category increases significantly (e.g. E14-E20: 22.9% to 59.2% p<0.001 and E16-E20 31.3% to 59.2% p<0.001) and an increase in the +/− 10⁰- 20⁰ group (E14-E16: 19.2% to 23.4% p<0.05). Additionally there is an accompanying reduction in the proportion of fibres in the third and fourth categories, +/−20° - 30° (e.g. E14-E20 14.8% to 8.2% p<0.01, E16-E20 15% to 8.2% p<0.01) and +/− 30⁰- 40⁰ e.g. E14-E20 10.5% to 3.7% p<0.001, E16-E20 9.8% to 3.7% p<0.001). This shift is demonstrated by the proportion of fibres represented within +/− 40⁰ of the longitudinal axis, increasing from 67.7% at E14, 79.5% at E16, to 94.2% at E20. In addition, the birefringence pattern shows a transition in the colours of the fibres, with the majority of fibres at E14 appearing green or yellow indicating young to intermediate maturity and an obvious transition to orange and red (indicative of more mature) present in E16 and E20, with predominantly red fibres at E20, seen across biological specimens in supplementary figure 2.

**Figure 5:**
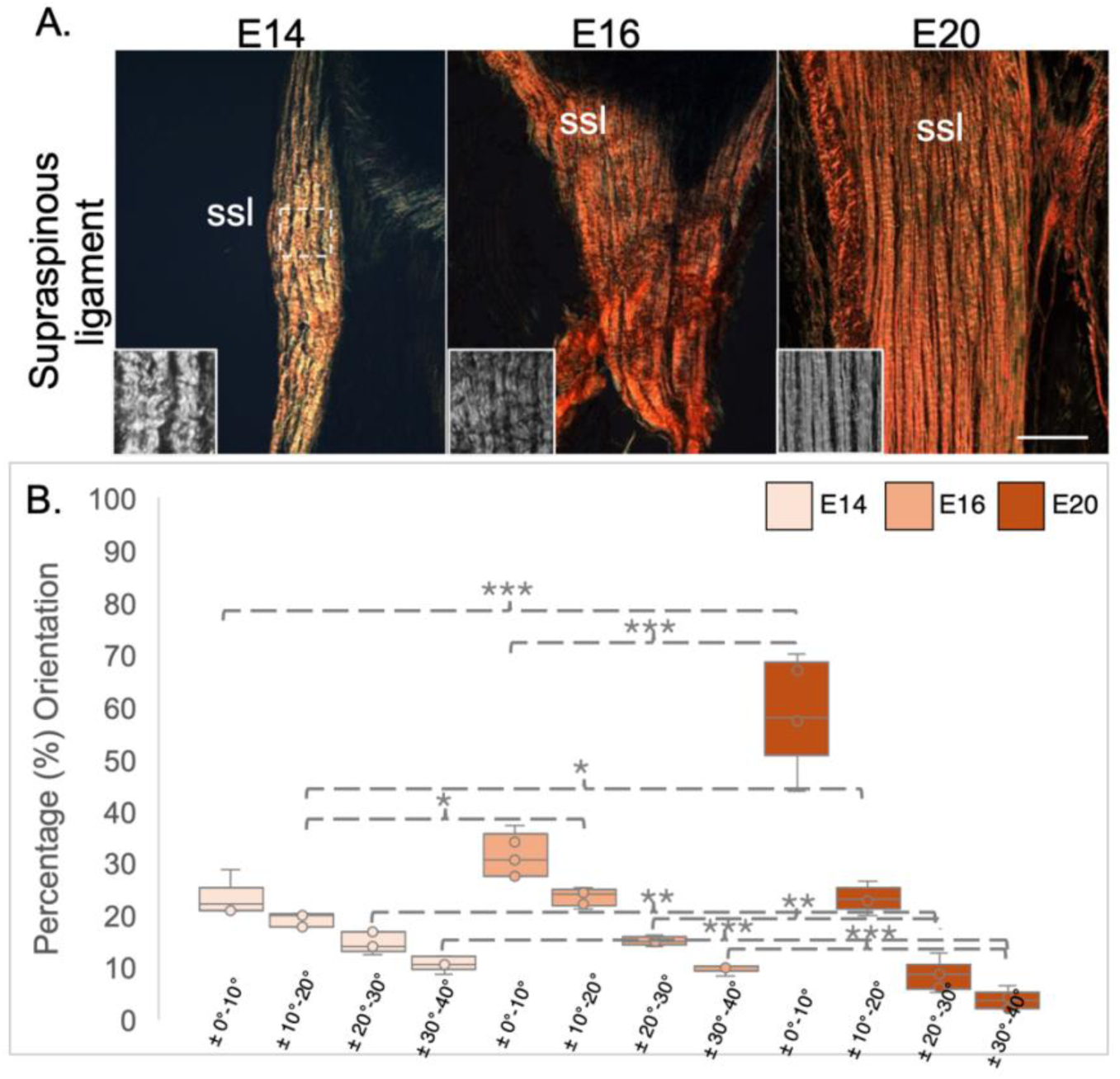
Collagen fibres become more aligned in dorsal spinal ligaments across development in the embryonic chick. (A) Collagen fibre bundles in supraspinous ligaments visualised by picrosirius red staining and polarised light microscopy (PLM) in longitudinal sections of supraspinous ligaments at E14, E16 and E20. Inserts represent regions of interest cropped and greyscaled during fibre orientation analysis. Vertical longitudinal view shown. (B) The box plot represents the proportion of fibres quantified following use of the OrientationJ plugin in ImageJ to analyse birefringence patterns, resulting in fibres plotted within four orientation groupings with respect to the spines long axis; +/− 0° - 10°, +/−10° - 20°, +/− 20° - 30°, +/− 30° - 40° (measurements taken from n=5 biological replicates for each embryonic stage). Scale bar 100µm. *p≤0.05, **p≤0.01, ***p≤0.001

### 3.4 Supraspinous ligament cells reduce in number and nuclei become smaller and more circular across development

Cell density in the dorsal supraspinous ligament was found to decrease across development, through haematoxylin and eosin staining with greater numbers of nuclei per 10,000μm^2^ at E14 and E16 (460±29.2 and 388±25.9) than at E20 (233±12.3, p=0.0000953 and p=0.00194, respectively)(Figure 6A and D) (Supplementary figure 3). Cell density is an indicator of maturity in connective tissues such as ligaments and tendons, with lower cell density indicating more mature tissues. This shows that across this timeframe these ligaments are becoming more mature, supporting the findings of section 3.3. There is a qualitative organisation of the nuclei across development, with an apparent alignment of cells in columnar channels at E20 (supplementary figure 3). Nuclei were found to be smaller at E20 (12.4±0.31μm^2^) than at E14 and E16 (15.4±0.4μm^2^, p= 0.00000000188 and 15.1±0.31μm^2^, p= 0.0000000000214) (Figure 6F) and the shape of the nuclei becoming more circular at later stages of development, quantified through aspect ratio, comparing E14 (2.53±0.05) to E16 and E20 (2.33±0.04, p=0.00492 and 2.37±0.045, p=0.04094) (Figure 6G). Lower aspect ratio values indicate that nuclei are becoming more circular, though they are still over twice as long as they are wide by E20.

**Figure 6:**
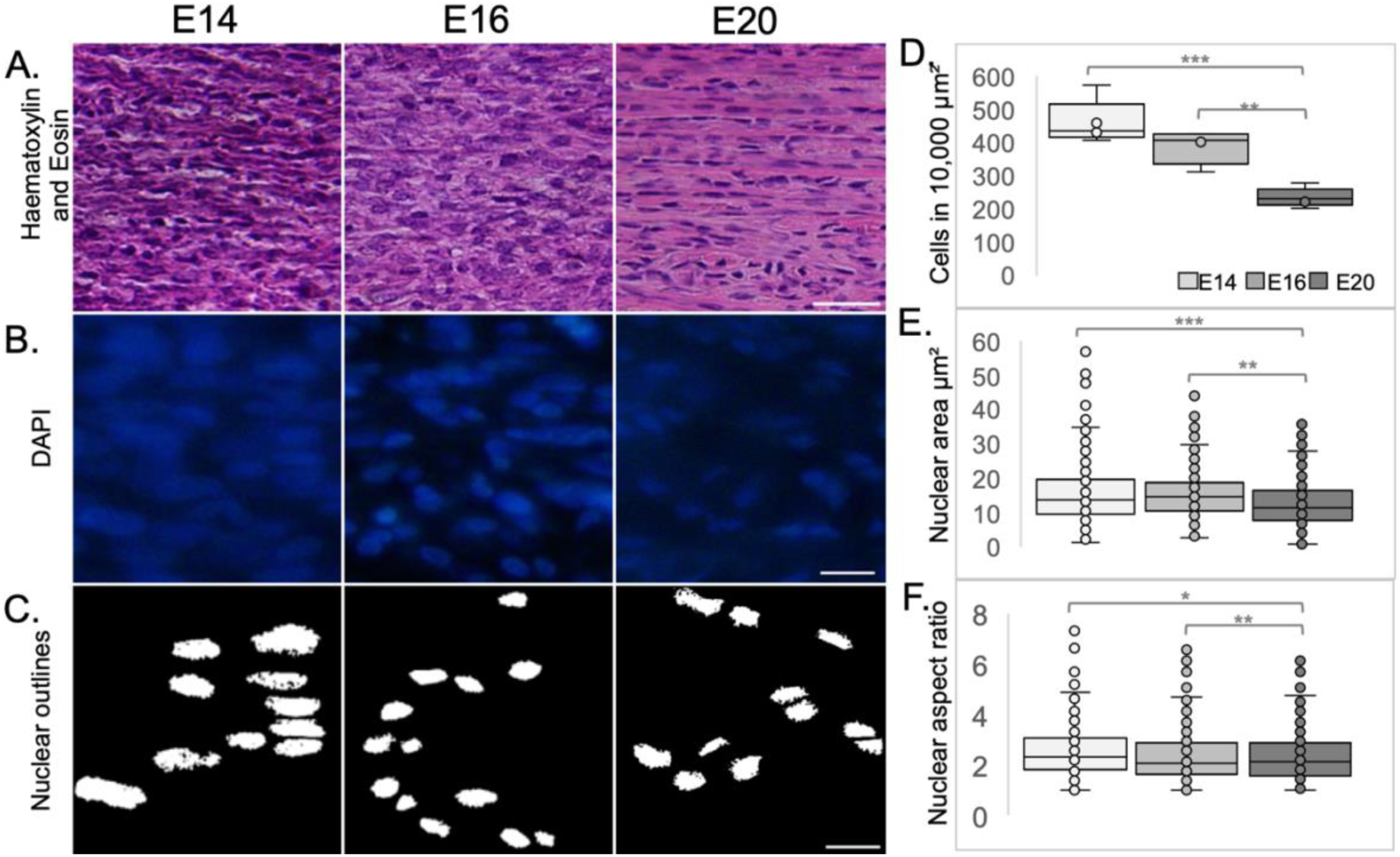
Cell density decreases and nuclei become smaller and more circular in the dorsal supraspinous ligament of the spine across development. (A) Dorsal supraspinous ligaments stained with haematoxylin (darker purple, stains cell nuclei) and eosin (lighter purple/pink, stains cytoplasm) across development. (B) Representative nuclei stained with DAPI and corresponding (C) thresholded image for nuclear analysis. (D) Cell density in 10,000μm² regions of interest (ROI) decreases across development, with (E) nuclear area decreasing and (F) nuclear aspect ratio (length/width) decreasing across development. Aspect ratio is a representative measurement for circularity. Scale bars: (A) 25μm (B&C) 10μm. *p≤0.05, **p≤0.01, ***p≤0.001.

## 4 Discussion

To better understand the developmental progression of spinal ligaments, as a first step toward understanding if and how these tissues contribute toward spinal abnormalities, such as scoliosis, this study provides the first microanatomical characterisation of developing spinal ligaments using the embryonic chick model. We histologically profile changes *in vivo* across the late stages of embryonic (E) E14 - E20 development in the chick spine. We identify six collagen-rich ligamentous structures in the embryonic chick spine, homologous to those in the mammalian spine, that interact directly with cartilaginous vertebral elements. To better understand tissue, cellular, and molecular changes that take place during spinal ligament maturation we describe the morphological changes that occur across late stages of development. With a focus on the thoracic spine we show that collagen is deposited in bundles that increase in size and area, with increasingly more mature and aligned collagen fibres across development. Molecular investigations show a maintenance of ECM associated gene expression. We demonstrate that cell density decreases and nuclei become less elongated and get smaller in the dorsal supraspinous ligament across E14 to E20.

Considerations as to the most appropriate model systems for studying the spine are required. Common developmental model systems used include the mouse and zebrafish (Busse et al., 2020), each offering benefit for genetic investigations amid different considerations (Terhune et al., 2022). One important consideration is the biomechanical environment. Both the mouse, quadrupedal horizontal spinal curvature, and the zebrafish, limbless horizontal spinal orientation, have spinal anatomies akin to their functional requirements. However, in order to be able to compare models most appropriately to the human spine we must carefully consider biomechanical factors. One important component of the musculoskeletal system that provides biomechanical stabilisation of the spine are ligaments, so to consider these as a functional unit, it is important to qualitatively assess these in a system that too has an upright structure. The spinal ligament morphology in the chick was observed to be analogous to that of the human, with intersegmental ligaments: anterior longitudinal, posterior longitudinal, supraspinous ligament being present, along with intrasegmental ligaments: ligamentum flavum, intervertebral and intertransverse ligaments being identified (Butt et al., 2015a). Intertransverse ligaments have been shown to be present in avians and humans, but not in quadrupeds (Jiang et al., 1995). The presence of the intertransverse ligament may be an adaptation to bipedalism, though this has not been tested by investigating the ligaments of other bipedal organisms, such as kangaroos. The presence of this intertransverse ligament, absent in other animals, may suggest a biomechanical similarity between both human and avian spines. Some aspects of spinal morphology in the chick are different to that of the human, as avians do not have intervertebral discs (Bruggeman et al., 2012), which account for up to 20–25 % of the height of the vertebral column in humans (Butt et al., 2015a). No involution of the notochord takes place in avians, and no nucleus pulposus is present (Bruggeman et al., 2012). Chick spines also have different numbers of vertebrae, having 14 cervical vertebrae and 7 thoracic vertebrae, while human spines have 7 cervical vertebrae and 12 thoracic vertebrae (Bui and Larsson, 2020; Butt et al., 2015a). Thoracic vertebrae 2 to 7 fuse during development in the chick, beginning at day 11 (Bui and Larsson, 2020). In spite of differences, the similarities in posture and the identification of homologous spinal ligaments in the chick suggest it is another appropriate model for experimentally investigating the spine. Collagen fibres run parallel to the long axis of tendons and ligaments, providing strength and mechanical stability (Birk et al., 1995; Birk et al., 1991; Kato and Silver, 1990), though not all ligaments show high levels of collagen alignment, even at maturity (Stender et al., 2018). Here, we show that fibre alignment increases across development in the embryonic chick spine, perhaps suggesting that these ligaments would show greater resistance to deformation (Szczesny et al., 2012) and providing more structural stability as development proceeds. Corresponding to the observed alignment of collagen fibres and nuclear changes observed in embryonic tendons (Rolfe et al., 2024) collagen fibres in spinal ligaments show similar alignment across development, with the mean percentage of fibres between +/− 0-20 degrees of straight as 82% in the supraspinous ligament, compared to 84% in the embryonic limb tendon at E20 (Rolfe et al., 2024). Fibre alignment has been shown to increase with maturity in chick tendons (Birk et al., 1995), along with an increase in fibre length and accompanying increase in strength (Peterson et al., 2024; Peterson et al., 2021).

Tendons and ligaments are both hypocellular connective tissues, with ligaments observed to have a greater cell density than tendons (Amiel et al., 1984). Connective tissues cellular density decreases with time, and our data show similar decreases in density as those observed in the embryonic chick limb tendon and jaw (Korntner et al., 2022; Rolfe et al., 2024). Nuclei of spinal ligaments, though showing increasing circularity, remain more elongated when compared to connective tissues in the limb at E20 (Rolfe et al., 2024), while nuclei in connective tissues of the jaw have been shown to become more elongated with time (Korntner et al., 2022). This indicates that nuclear shape in connective tissues may show site specificity, a finding supported by observations of different ligaments in rabbits showing different cellular shapes at full maturity (Amiel et al., 1984). In addition the cellular organisation in the supraspinous ligament demonstrates organisation into aligned columns at E20, similar to those observed in embryonic tendons (Rolfe et al., 2024). Our data suggest that there are similarities in the structural organisation of spinal ligaments and limb tendons across development, with fibres becoming more aligned and the density of cells reducing, yet the shape of the nuclei demonstrating differences. This is consistent with previous reports on ligament and tendon development, showing that collagen alignment and fibril diameter correlate with function and mechanical strength (Carballo et al., 2018). Additionally, future studies on spinal ligament biomechanics must be performed to uncover the importance of this structure for the tissues tensile strength.

To explore the molecular changes that accompany embryonic ligament development and maturation we profiled expression changes in the dorsal supraspinous ligament showing a reduction in the transcriptional regulator *Scx* across development, with reductions in collagen component genes and maintenance of extracellular matrix associated genes *fibrilin1* and *decorin*. *Scx* is a helix-loop-helix transcription factor which is the earliest known marker progenitor cell populations in early development of ligaments and tendons (Nakamichi and Asahara, 2021; Schweitzer et al., 2001). *Scx* also promotes tenomodulin expression later in development, with tenomodulin being a type II transmembrane glycoprotein expressed in ligaments, tendons, and eyes and is a regulator of tenocyte proliferation and is involved in collagen fibril maturation (Docheva et al., 2005). *Scx* demonstrates a similar decrease in expression across development in embryonic spinal ligaments as in embryonic limb tendons (Rolfe et al., 2024) corresponding to the maturation of the tissue. *Fibrillin1* encodes fibrillin 1, a protein which makes up some of the elastic fibers found in the extracellular matrix of ligaments and tendons, along with collagen, fibrillin 2, and elastin (Giusti and Pepe, 2016). Maintenance of fibrillin1 indicates that deposition of fibrillin1 is still occurring at the latest stage of development profiled here. *Decorin* encodes a small leucine-rich proteoglycan (SLRP) which, in ligaments and tendons, makes up part of the extracellular matrix by binding to collagen and playing a role in fibril organisation and structure (Reed and Iozzo, 2002). The maintenance observed in chick embryonic spinal ligaments indicates that this structural organisation is continual, with the importance of this ECM molecule reinforced by its experimentally depletion importance in healing ligaments (Nakamura et al., 2000). While it is known that tendons and ligaments are vital to the transmission of forces and stabilisation of the musculoskeletal system (Bobzin et al., 2021), this study adds new information about the structural hallmarks of spinal ligaments, a ligamentous tissue not previously profiled, adding information about tissue, cellular and molecular characteristics.

Justification for the investigation of spinal ligaments to understand spinal defects comes from reports that defects in ligamentous tissues are observed in clinical presentations of scoliosis; including reduced fibre densities and distributions (Hadley-Miller et al., 1994), reduced ligament stiffness or ligament laxity in scoliosis patients compared to healthy individuals (Crijns et al., 2017; Zhang et al., 2022). In addition, connective tissue disorders (Marfan’s syndrome) commonly display scoliosis (Chotigavanichaya et al., 2022; Glard et al., 2008). Recent experimental evidence implicates a mechanosensitive (Musa et al., 2019) cell surface receptor to be required in spinal ligaments for spinal alignment in a mouse model of AIS (Liu et al., 2021b), and deficiencies in mechanically responsive ion channels *Piezo2* and *Asic2* have been shown to lead to scoliosis (Assaraf et al., 2020; Bornstein et al., 2023; Wu et al., 2020). This highlights the importance of biomechanical considerations when investigating the spine. Combining the advantages of the chick embryonic experimental model that allows for movement manipulation, with this insight into spinal ligaments, will allow for future investigations of the role mechanical stimulation plays in spinal ligament tissue patterning.

Taken together this study has established that the embryonic chick is an appropriate experimental model to study spinal ligaments, the morphological structures responsible for spinal stability. It details the structural hallmarks and provides the first insight into mechanistic changes that occur in these spinal tissues across the final stages of chick embryonic development. This will allow for further investigation of the molecular and biomechanical mechanisms required for ligament development.

## Author contributions

SH performed experiments, analysed data, drafted the manuscript.

ETB performed experiments, reviewed the manuscript.

RR performed experiments, conceptualised experiments, analysed data, drafted the manuscript.

## Acknowledgements

The authors are grateful to Prof Paula Murphy for her expert advice with respect to the conceptualisation of this work and the experimental components involved and to Lauren Sliney for her for technical support during.

## Funding Information

This work was supported by an Enterprise Ireland Horizon Europe Co-ordinator proposal preparation support scheme grant (EC20231473), a Higher Education research grant (2024) and with thanks to support of the Fondation Yves Cotrel pour la Recherche en Pathologie Rachidienne - Institut de France”.

## Conflict of Interest Statement

All authors of this manuscript declare that they have no conflicts of interest to report. We confirm that all authors were fully involved in the study and preparation of the manuscript, and that the material within has not been submitted for publication elsewhere.

## Supplementary Material

The Supplementary Material for this article can be found on corresponding document.

## Notes

### Competing Interest Statement

The authors have declared no competing interest.

