## Supplementary Figures for "Characterisation of spinal ligaments in the embryonic chick"

Hennigan et al., In Preparation for Journal of Anatomy April 2025

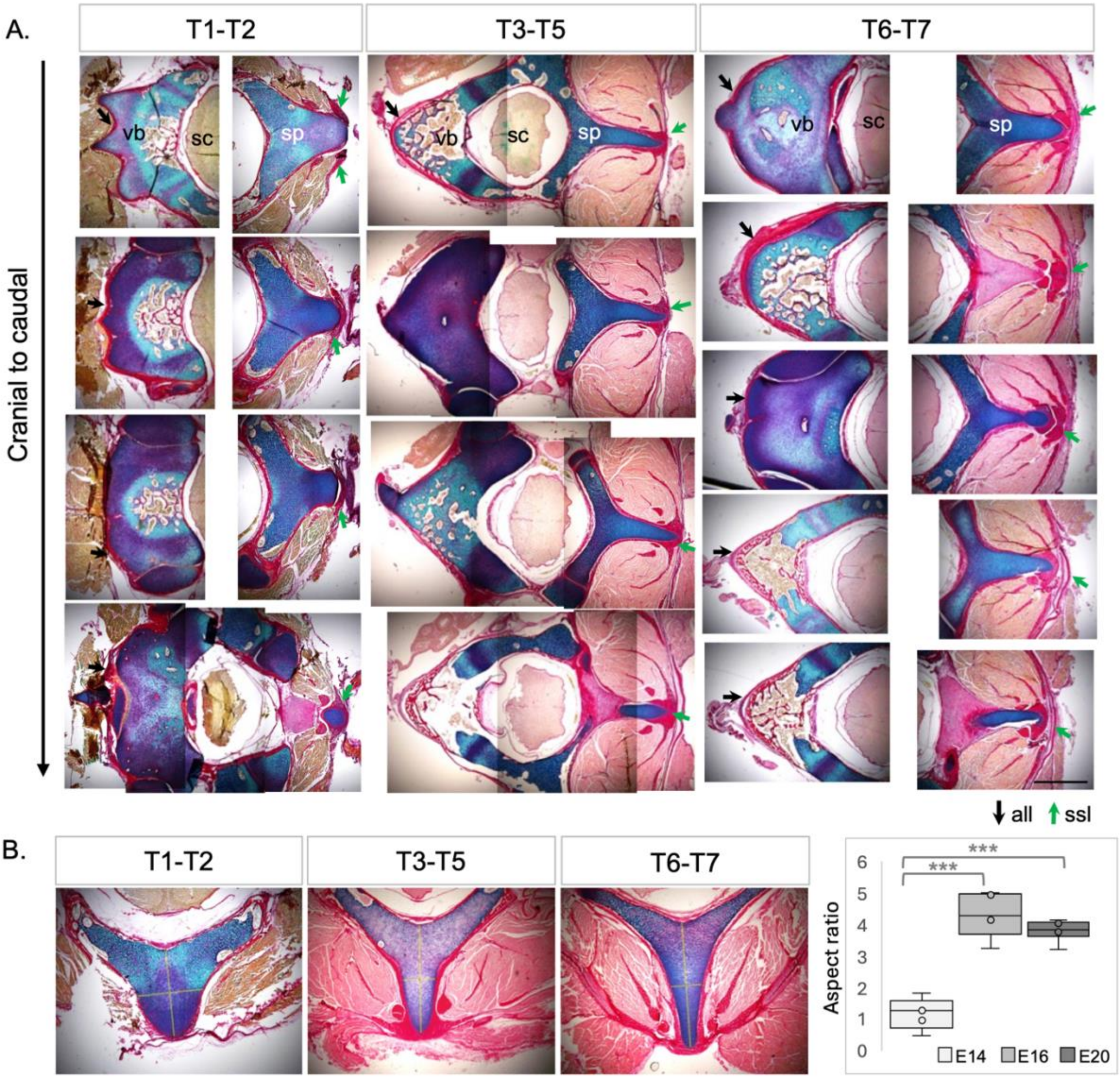


**Supplementary Figure 1**: The changing morphological shape of the vertebrae and the elaborate patterns of branching of the supraspinous ligament (dorsal) within the thoracic region are shown for each vertebral level, T1-T2, T3-T5 and T6-T7. (A) Each region shows the same biological specimen with four vertebrae imaged that run from a cranial to caudal direction, representing an approximate distance of 1,800μm (for T1-T2 and T3-T5) and 2,400μm (T6-T7) from top to bottom. (B) Box plot showing spinous process elongation in the aspect ratio representing the length and width of the cartilaginous spinous process across sub-regions of the thoracic vertebrae at E20. vb; vertebral body, sp; spinous process, sc; spinal cord, ssl; supraspinous ligament, all;. anterior longitudinal ligament. Scale bar 1000μm.

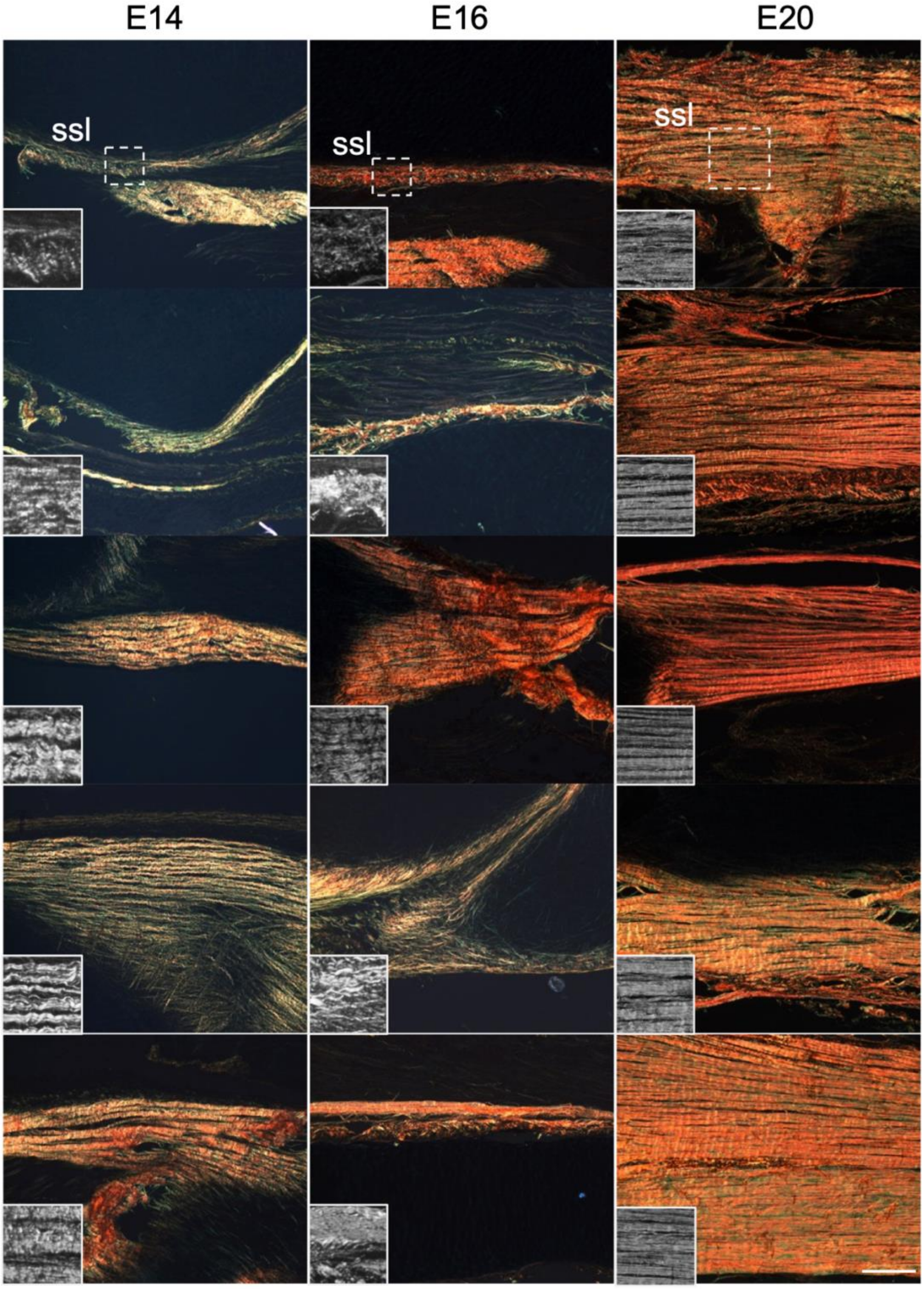


**Supplementary figure 2**: Collagen fibres become more aligned in spinal ligaments across development in the embryonic chick. Individual images represent different biological replicates (x15 in total) quantified for collagen fibre alignment across development (E14- E20), using OrientationJ analysis. Images showing longitudinal sections, orientated horizontal from cranial to caudal presenting the birefringence colour change indicating increasing fibre maturity. Fibres at E14 being green and yellow, to the emergence of orange fibres at E16 to fibres being predominantly red or orange at E20. Inserts represent regions of interest cropped and greyscaled during fibre orientation analysis. Scale bar 100µm.

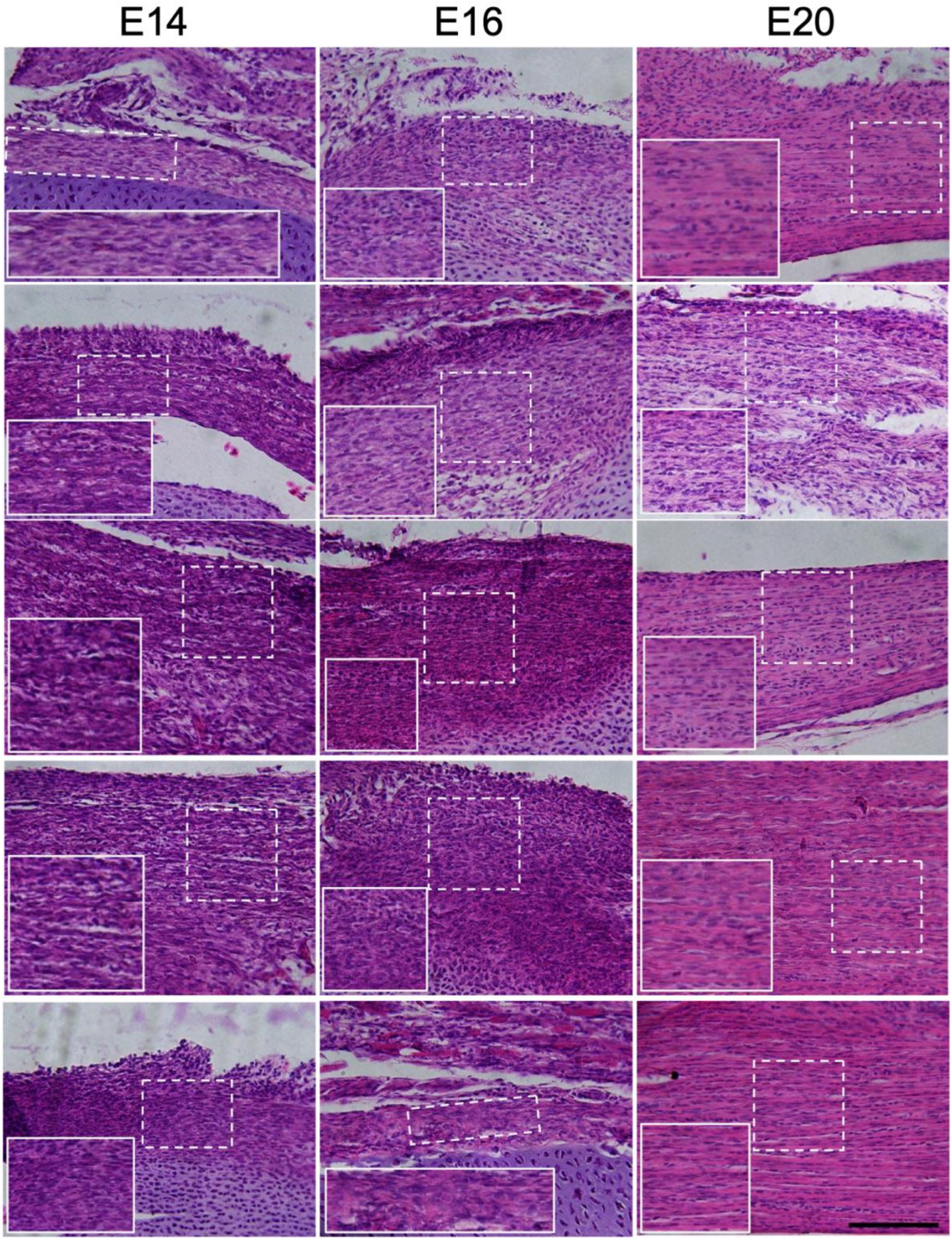


**Supplementary figure 3:** Cell density decreases across development. Images represent longitudinal views of dorsal supraspinous ligaments, stained with haematoxylin (darker purple, stains cell nuclei) and eosin (lighter purple/pink, stains cytoplasm). Five biological replicates are represented for E14, E16, E20 each. Images were cropped to regions of interest shown in bottom left of each image, from origin shown in dashed boxes. Scale bar 100µm.
